# Induction of the BIMS Splice Variant Sensitizes Proliferating NK Cells to IL-15 Withdrawal

**DOI:** 10.1101/392985

**Authors:** Benedikt Jacobs, Aline Pfefferle, Dennis Clement, Jodie P. Goodridge, Michelle L. Saetersmoen, Susanne Lorenz, Merete Thune Wiiger, Karl-Johan Malmberg

## Abstract

Adoptive transfer of allogeneic NK cells holds great promise for cancer immunotherapy. There is a variety of protocols to expand NK cells *in vitro*, most of which are based on stimulation with cytokines alone or in combination with feeder cells. Although IL-15 is essential for NK cell homeostasis *in vivo*, it is commonly used at supra-physiological levels to induce NK cell proliferation *in vitro*. As a result, adoptive transfer of such IL-15 addicted NK cells is associated with cellular stress due to sudden cytokine withdrawal. Here, we describe a dose-dependent addiction to IL-15 during *in vitro* expansion, leading to caspase-3 activation and profound cell death upon IL-15 withdrawal. NK cell addiction to IL-15 was tightly linked to the BCL-2/BIM ratio, which rapidly dropped during IL-15 withdrawal. Furthermore, we observed a proliferation-dependent induction of BIM short (BIM S), a highly pro-apoptotic splice variant of BIM, in IL-15 activated NK cells. These findings shed new light on the molecular mechanisms involved in NK cell apoptosis following cytokine withdrawal and may guide future NK cell priming strategies in a cell therapy setting.

## Introduction

Natural killer (NK) cells are commonly referred to as innate lymphocytes that display strong cytolytic potential in the absence of prior sensitization (1). However, the acquisition of functional potential is dependent on exposure to homeostatic cytokines as illustrated by the priming of naïve NK cells through trans-presentation of IL-15 by dendritic cells (2, 3). Cytokine stimulation rapidly induces enhanced effector function in NK cells (4-6), suggesting that all NK cells have an intrinsic ability to reach a cytolytic phenotype given sufficient stimulation. This holds true even from the early stages of differentiation in the complete absence of both positive and negative receptor input (7). The most important homeostatic cytokine during NK cell development and for survival in the periphery is IL-15 (8-10). IL-15 induces robust NK cell proliferation, NK cell differentiation, up-regulation of granzyme B and increases effector responses, including cytokine production and degranulation (5, 7, 11, 12).

The multi-faceted role that IL-15 plays in NK cell homeostasis, activation and differentiation has been linked to the induction of different downstream pathways dependent on the cytokine concentration (13). Whereas low doses of IL-15 induced the STAT5 pathway, ensuring development and survival, high doses induced the mammalian target of rapamycin (mTOR). mTOR activation increased the cell’s metabolic activity, switching from primarily utilizing oxidative phosphorylation to glycolysis, a process termed metabolic reprogramming (14). This switch in energy source was necessary to maintain effector potential.

In the clinical setting, cytokines are used to prime NK cells for adoptive cell therapy to enhance cytolytic potential and to ensure sufficient numbers through the induction of proliferation. Supra-physiological levels of various cytokines, including IL-2, IL-15, IL-12 and IL-18 and combinations thereof, are often used during *in vitro* expansion (15). However, due to severe side effects, these high doses of cytokines cannot be administrated to patients to further support expansion *in vivo* (16-19). Consequently, NK cells experience a strong reduction in cytokine concentration upon adoptive transfer, which can severely affect their functional potential, survival and long-term engraftment (20). Although we largely lack immunobiological correlates of good outcomes in NK cell trials, a common denominator for success appears to be *in vivo* persistence and expansion of donor-derived NK cells (21). In high-risk or refractory acute myeloid leukemia (AML) patients treated with IL-2 activated, haploidentical NK cells, persistence and expansion of donor NK cells were associated with higher rates of complete remission and increased progression-free survival (22, 23). Furthermore, in refractory non-Hodgkin-lymphoma patients, the response to adoptive NK cell therapy has been linked to levels of endogenous IL-15 at the day of NK cell infusion (19). The need for *in vivo* expansion of adoptively transferred NK cells highlights the importance of understanding the mechanisms regulating NK cell homeostasis and the cell fate after sudden deprivation of high cytokine concentrations.

Recently Mao Y. et al., reported that NK cells primed with a short pulse of IL-15 were better at surviving and sustaining cytolytic activity upon cytokine withdrawal compared to IL-2 primed NK cells (20). IL-15 has been shown to enhance NK cell survival through the complex interaction of BCL-2 family members (24). Indeed, the improved survival in IL-15 primed NK cells compared to those primed with IL-2 was linked to STAT5-dependent up-regulation of BCL-2, while the sustained cytolytic activity upon cytokine withdrawal was attributed to enhanced mTOR signaling upon IL-15 priming. However, it remains unclear how prolonged exposure to IL-15 and induction of proliferation influence the expression and dynamic balance of pro-and antiapoptotic molecules in discrete NK cell subsets. The BCL-2 family members can be divided into 3 groups, the anti-apoptotic, pro-apoptotic effector and BH3-only proteins, which together effectively control the intrinsic apoptosis pathway. The anti-apoptotic BCL-2 family members (MCL-1, A1, BCL-XL and BCL-2 itself) inhibit apoptosis by binding pro-apoptotic proteins, like the BH 3-only proteins BIM, NOXA or PUMA (25). These pro-apoptotic BH3-only proteins can either directly or indirectly (by reducing the availability of anti-apoptotic proteins) activate the pro-apoptotic effector proteins BAX and BAK (26). Once these two major players of the intrinsic apoptosis pathway are activated, they oligomerize and permeabilize the outer mitochondrial membrane leading to the release of cytochrome c into the cytoplasm and subsequently to the activation of executor caspases (27).

Here, we investigated the dynamic regulation of pro-and anti-apoptotic molecules during IL-15 induced proliferation/activation and subsequent withdrawal. Our data reveal a dose-dependent addiction to IL-15 that correlated with the degree of proliferation achieved during the *in vitro* expansion phase. This addiction resulted in increased susceptibility to apoptosis upon sudden IL-15 withdrawal and correlated with an altered expression of anti-and pro-apoptotic BCL-2 family members. In particular, we found that the highly pro-apoptotic BIM short (BIM S) splice variant accumulated in NK cells exposed to high concentrations of IL-15, suggesting a mechanism by which these cells are prone to rapid apoptosis. These results shed new light on the molecular mechanism of cytokine addiction in human NK cells following IL-15 driven expansion.

## Materials and Methods

### Reagents and cell lines

Antibodies were purchased for CD57 FITC/ PE, Streptavidin BV785, BCL-2 AF647 from BioLegend, for CD3 V500, CD14 V500, CD19 V500, anti-IgM BV650, MCL-1 purified from BD Bioscience, for CD56 ECD from Beckman Coulter, for CD57 functional grade purified from eBioscience, for BIM AF647, BCL-XL AF647 from Cell Signaling Technologies and for NKG2A PE-Vio770, KIR2D biotin, KIR3DL1/2 biotin from Miltenyi Biotec. The MCL-1 purified antibody was labeled with an AF647 antibody labeling kit from Life Technologies accordingly to the kit’s instructions. The BCL-2 family member inhibitors ABT-199 and ABT-737 were purchased from Santa Cruz BioTechnologies and the MCL-1 inhibitor Maritoclax from Tocris. Pacific Orange and Blue Succinimidyl Ester were bought from Thermo Fisher Scientific. The pan-caspase (Z-VAD-FMK) and caspase-8 (Z-IETD-FMK) inhibitors were purchased from R&D Systems. K562 cell line from ATCC was cultured in RPMI 1640 media with antibiotics (penicillin/streptomycin; Sigma) and 10% heat-inactivated fetal calf serum (Sigma) at 37°C.

### NK cell isolation and culture

Buffy coats from random healthy blood donors were purchased from the Oslo University Hospital Blood bank with donor informed consent. Using density gravity centrifugation (Lymphoprep; Axis-Shield) and fretted spin tubes (Sepmate, Stemcell Technologies) peripheral blood mononuclear cells (PBMCs) were isolated and used for NK cell isolation. NK cells were isolated from PBMCs using a NK cell isolation kit and an AutoMACS Pro Separator (Miltenyi Biotec). Freshly isolated NK cells were labeled with the CellTrace^™^ Violet or CFSE^™^ dye for cell proliferation analysis according to the kit’s instructions (Molecular Probes). CTV-/CFSE-labeled NK cells were culture in RMPI 1640 media (Sigma) with antibiotics (penicillin/streptomycin; Sigma) and 10% human, heat-inactivated AB serum (Trina Bioreaktives) plus 10 or 1 ng/ml IL-15 (Miltenyi Biotec) for 6 days at 37°C. On day 2 and 4 the medium was replaced with fresh medium and IL-15. After 6 days cells were harvested, washed and counted. Cells were either used for analysis or incubated further for up to 48h in medium with or without the prior IL-15 dose at 37°C.

### Flow Cytometry staining

Freshly isolated or IL-15 treated NK cells were stained in staining buffer (PBS + 2% fetal calf serum (FCS) + 2 mM EDTA) with various antibody combinations and a dead-cell marker (LIVE/DEAD® Fixable Near-IR or Aqua Dead cell stain kit; Life Technologies). Cells were fixed afterwards in 2-4% paraformaldehyde (PFA) and either directly analyzed at a LSRII flow cytometer instrument (Becton Dickinson) or stained for intracellular proteins after permeabilization with 100% methanol at –20°C.

Data was analyzed using the FlowJo V10.0.8 software (TreeStar). The gating strategy is illustrated in supplementary figure 1A+B.

**Figure 1.**
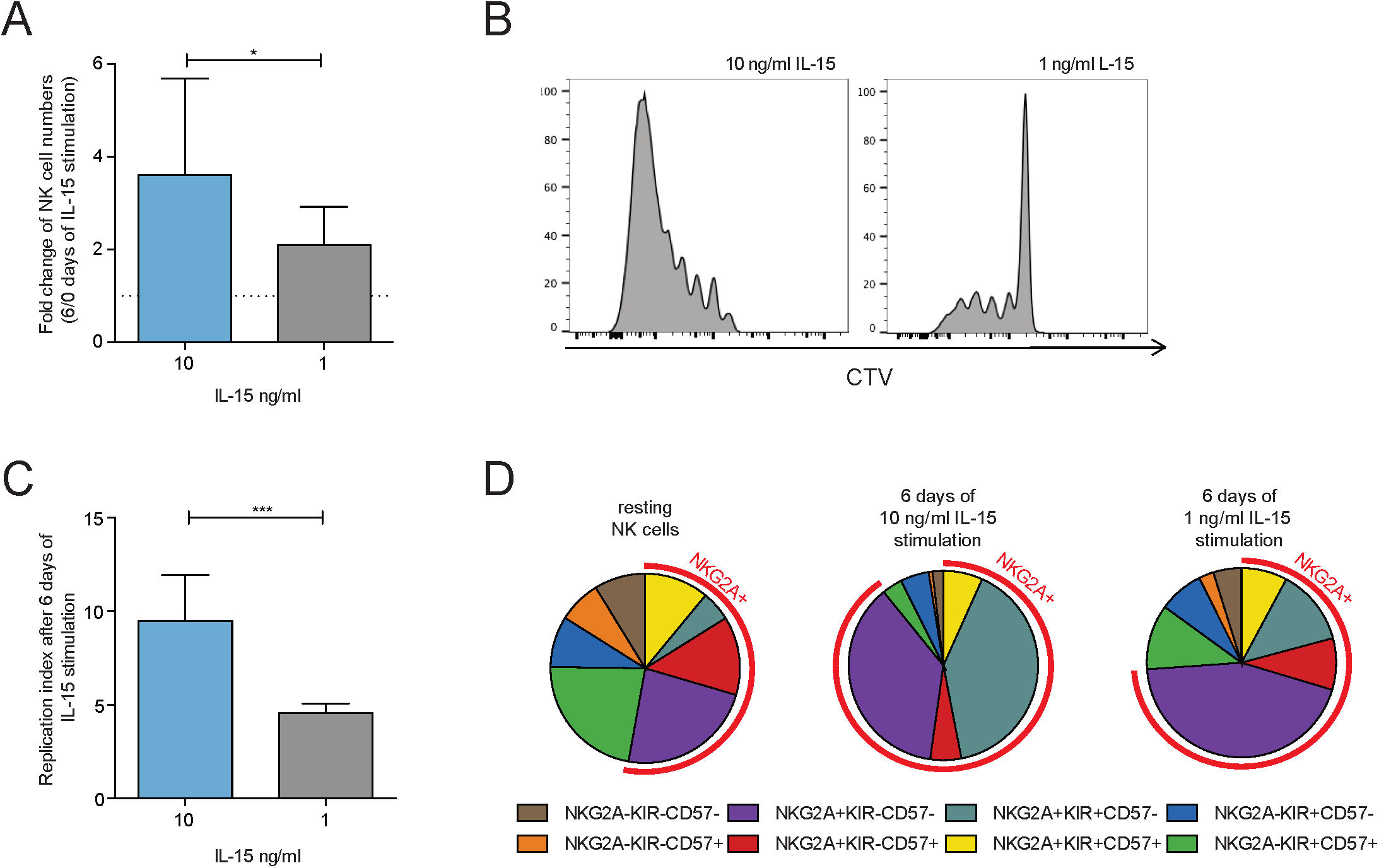
**A)** The fold increase in cell number of freshly isolated NK cell between the start (day 0) and after 6 days of high-(10 ng/ml) or low-dose (1 ng/ml) IL-15 treatment (n ≥ 6). **B)** Representative histogram of the CTV-dilution in NK cells treated for 6 days with high-or low-dose IL-15. **C)** Based on the NK cell numbers and their CTV-dilution after 6 days of IL-15 treatment (high-vs. low-dose), a *replication index* (fold expansion of dividing cells) was calculated for both treatment groups (n = 14). **D)** Pie charts of the distribution of different NK cell subsets based on their expression of NKG2A, KIR and CD57 after isolation from peripheral blood and after 6 days of high-or low-dose IL-15 treatment. Significance were calculated using a Wilcoxon test (p-values: * <0,05; ***<0,001).

### Barcoding and BCL-2 family member staining

In order to improved staining quality of intracellular proteins and to reduce the amount of antibodies used, differently treated NK cells were labeled after methanol permeabilization with a unique dilution combination of the two dyes pacific blue and orange (Life Technologies). Afterwards cells were collected together and stained for various surface markers and intracellular BCL-2 family member proteins. An example of the gating strategy for IL-15 treated NK cells, which have been stained with BCL-2 is illustrated in supplementary figure 1C. In order to compare the expression intensity of BCL-2 family members between different NK cell donors across multiple experiments, the MFI values were normalized to NK cells isolated from a reference donor included in each fluorescent barcoding experiment.

### Analyzing NK cell proliferation

To analyze the increase in NK cell numbers upon IL-15 treatment, their numbers were measured at the start of the IL-15 treatment and after 6 days. FlowCount^™^ Fluorospheres (BeckmanCoulter) were used to calculate the NK cell numbers per µl and a ratio between the NK cell numbers on day 6 and at the start of the IL-15 treatment was calculated. The analysis for fold change in NK cell numbers upon continuous IL-15 treatment after 6 days or IL-15 withdrawal was done accordingly.

To analyze the degree of cell proliferation during the 6 day IL-15 treatment, CTV-/CFSE-dilution was measured on day 6 by flow cytometry. Afterwards the replication index (fold expansion over culture time of proliferating cells; 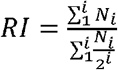 with i=generation number and N_i_ = number of events in generation i) was calculated (28).

### Measuring Caspase-3 activity

NK cells were cultured for a duration of 6 days with 10 or 1 ng/ml IL-15 followed by a varying period of continued culture (up to 48 hours) with or without the prior IL-15 dose. 1 hour before the end of the culture the caspase-3 inhibitor DEVD-FMK conjugated to FITC (Caspase-3 active FITC staining kit; Abcam) was added to the culture to label cells with active caspase-3 activity. Cells were harvested and stained for NK cell surface marker and analyzed by flow cytometry.

### Western Blotting

NK cells were harvested, washed twice with ice-cold PBS and lysed in 50-100 µl RIPA buffer plus protease/phosphatase inhibitors (Thermo Fisher Scientific) for 15’ on ice and then stored at-80°C. Protein lysates were thawed and the protein concentration was evaluated using a Pierce BCA protein assay kit (Thermo Fisher Scientific). Protein lysates were cooked at 90°C for 5’ with NuPage® LDS sample buffer (Thermo Fisher Scientific). 10 µg protein were loaded onto a NuPAGE® Novex® Bis-Tris 12% MiniGel (Life Technologies) and run for 45-60’ at 150 V. Afterwards, a wet blotting transfer was done onto a 0,2 µm PVDF membrane (Life Technologies) for 90’ at 4°C and 100 V. The membrane was blocked for 1h in TBST buffer (1x TBS + 0,1% TWEEN 20) plus 5% BSA and incubated with the BIM antibody (Clone: C34C5; 1:1000; Cell Signaling Technologies) overnight. Next day, the membrane was washed 3x with TBST buffer and incubated for 1 h at room temperature with an anti-rabbit HRP antibody (1:3000; Dako). The membrane was washed 3x with TBST buffer, visualized with a Pierce ECL western blotting substrate (Thermo Fisher Scientific) at a Chemidoc machine (BioRad) and acquired with the Image Quant software. The staining for Actin B was done accordingly. The analysis was performed using ImageJ (NIH).

### Peggy Sue^*™*^ instrument

To analyze BCL-2 family members in sorted NK cells, capillary electrophoresis and immunodetection of proteins were performed in the Simple Western system Peggy Sue^*™*^ using the 2-40 kDa size separation kit. Cells were lysed in RIPA buffer plus protease/phosphatase inhibitors (Thermo Fisher Scientific) for 15’ on ice and stored at-80°C. Protein lysates were thawed and the protein concentration was evaluated using a Pierce BCA protein assay kit (Thermo Fisher Scientific). For each sample 0.2 mg/ml protein lysates were used. Primary antibodies were diluted 1:50, and ready-made secondary antibodies were used from Protein Simple. Protein lysates and reagents were pipetted into a 384 well plate, centrifuged at 2000x *g* for 5’ and put into the Peggy Sue^™^ machine. Experimental set-up and data analysis were done using the Compass software (Protein Simple).

### RNA sequencing

Freshly isolated NK cells and 5 day IL-15 stimulated NK cells were sorted into three subsets (NKG2A^+^KIR^-^CD57^-^,NKG2A^-^KIR^+^CD57^-^,NKG2A^-^KIR^+^CD57^+^CD38^low^/NKG2C^+^) using a FacsAria (BD) at 4°C. Day 5 cells were further sorted into proliferating cells (generation 1) using CellTrace Violet staining. RNA was isolated and the library was prepared using the Illumina NeoPrep Library preparation system. NextSeq(Illumina) (single read, 75base pairs) was used for sequencing and Bowtie (version 2.0.5.0) and Tophat (version 2.0.6) were used to carry out the read alignment. Cufflinks (version 2.1.1) was used to estimate the transcript abundance. FKPM values from the individual subsets were pooled for a global analysis comparing baseline (day 0) to proliferating (day 5). The log2 fold change and adjusted p value were used for visualization in the volcano plot.

### Statistical analysis

For the comparison of single matched groups or populations a Wilcoxon test was used. A Wilcoxon signed rank test was performed when calculating the statistical significance of a given median to a hypothetical value. For comparing multiple matched groups with each other, a one-or two-way ANOVA test was done. Statistical significance: ns indicates not significant, * p<0.05; **p<0,01, ***p<0,001,****p<0,0001. Analysis was performed using the GraphPad Prism software.

## Results

### High-dose IL-15 skews NK cells towards a more immature phenotype

To study the molecular consequences of cytokine-withdrawal, primary human NK cells were stimulated with either high-(10 ng/ml) or low-dose (1 ng/ml) IL-15 for 6 days *in vitro*. After 6 days, NK cell numbers increased significantly in high-dose IL-15 treated cells (Figure 1A), reflected in a stronger CTV dilution and higher replication index than those treated with low-dose IL-15 (Figure 1B-C). NK cells can be divided into various subsets based on their expression of NKG2A, KIRs and CD57, which define their level of differentiation (6). Although treatment with both concentrations of IL-15 increased the fraction of NKG2A^+^ NK cells and decreased the fraction of the more differentiated CD57^+^ NK cells, this effect was more pronounced in high-dose IL-15 treated NK cells. Furthermore, high-dose IL-15 treatment resulted in a dramatic increase in the NKG2A^+^KIR^+^CD57^-^ subset fraction (Figure 1D). Together, these initial experiments established a platform for studying the effect of cytokine withdrawal in discrete NK cell subsets based on their degree of prior activation.

### Strongly proliferating NK cells are more susceptible to IL-15 withdrawal

To monitor the effect of IL-15 withdrawal after NK cell expansion, isolated NK cells were pretreated for 6 days with high-or low-dose IL-15. Subsequently, cells were harvested and put into culture again for 48 h with the either the same IL-15 concentration as before or with IL-15 being completely withdrawn (Figure 2A). The effect of continued IL-15 treatment versus IL-15 withdrawal was investigated by calculating the fold change in NK cell numbers before and after the additional 48 h incubation period for each treatment condition. As expected, continued IL-15 treatment for 48 h resulted in increased NK cell numbers, which were significantly higher if cells were pretreated with high-compared to low-dose IL-15. In contrast, NK cell numbers decreased in a dose-dependent manner 48 h after cytokine withdrawal (Figure 2B). Furthermore, we observed a significant correlation between the decrease in NK cell numbers upon IL-15 withdrawal and the replication index after the initial 6 days of IL-15 stimulation (Figure 2C).

**Figure 2.**
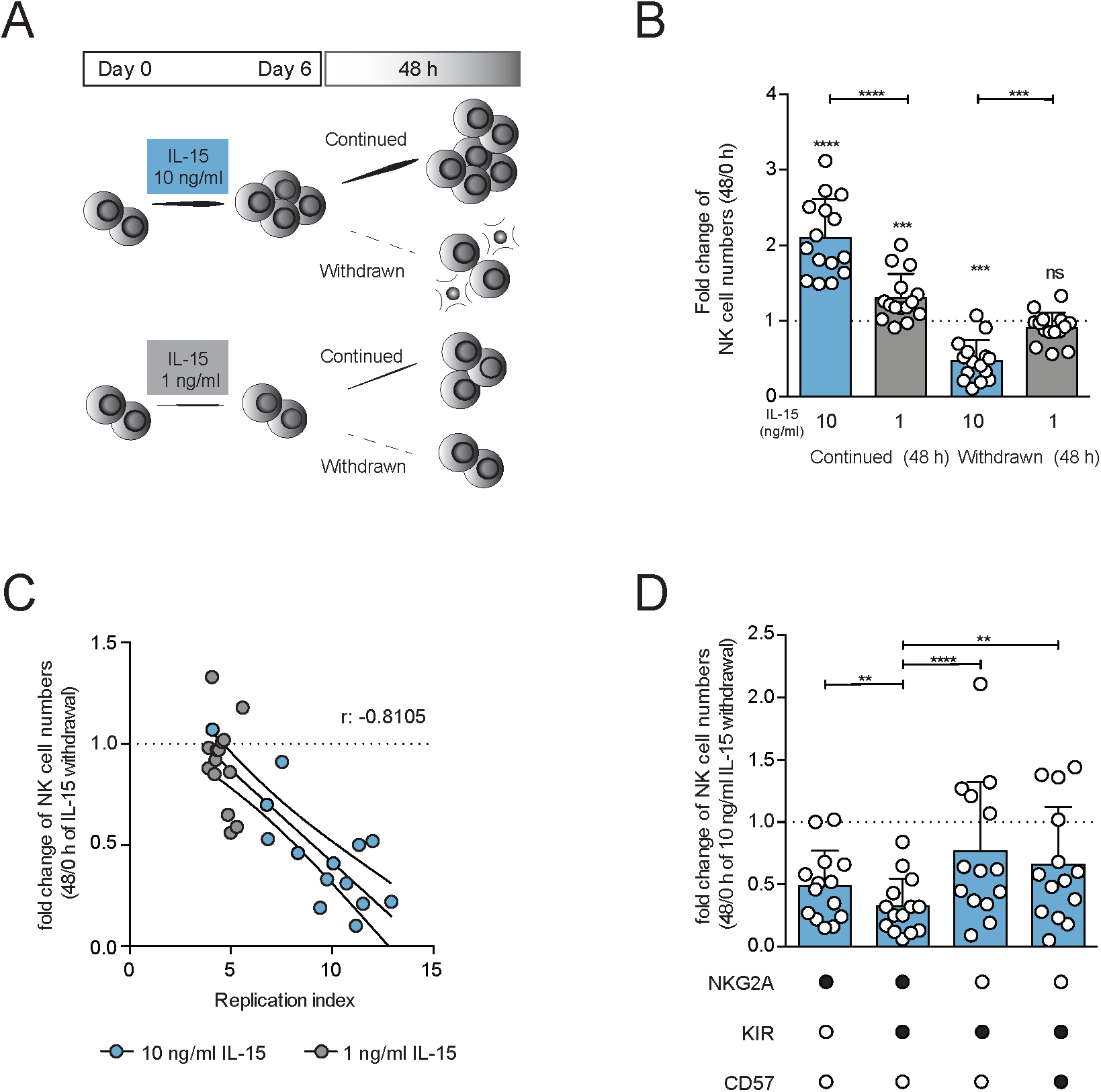
**A)** Illustration of *in vitro* NK cell treatment with IL-15 stimulation and withdrawal. NK cells were isolated from healthy donors and cultured with high-(10 ng/ml) or low-dose (1 ng/ml) IL-15 for 6 days. Cells were then harvested, washed, counted and put back into culture with (full line) or without (dashed line) the prior IL-15 dose for an additional 48 h. **B)** NK cells were treated for 6 days with high-(10 ng/ml) or low-dose (1 ng/ml) IL-15, harvested and put into fresh media with (+) or without (-) the prior IL-15 dose for 48 h. Fold change in NK cell numbers after the additional 48 h treatment (+/-IL-15) compared to numbers at day 6 (n = 15). Significance was calculated using a Wilcoxon signed rank test (stars alone), while a Wilcoxon test was used (stars above line) to calculate the significance between samples pretreated with high-or low-dose IL-15. **C)** A correlation between the decrease in NK cell numbers upon 48 h of IL-15 withdrawal and their replication index (fold expansion of dividing cells) after 6 days of high-(10 ng/ml) or low-dose (1 ng/ml) IL-15 treatment was calculated (n = 14). Correlation between fold change of NK cell numbers and their replication index was calculated using a spearman r test. **D**) The fold decrease in NK cell numbers after 48 h of IL-15 withdrawal was calculated for four discrete NK cell subsets, which had undergone prior treated of high-dose IL-15 for 6 days (n = 14). Significance of the fold change of NK cell numbers between different NK cell subsets was calculated using a Friedman test (p-values: ** <0,01; ***<0,001; ****<0,0001).

Next, we addressed whether NK cell differentiation influenced the susceptibility to IL-15 withdrawal. To this end, we analyzed the relative change in NK cell numbers of four discrete NK cell subsets at various stages of differentiation, following cytokine stimulation and withdrawal. In line with their differential intrinsic potential for proliferation (6), the decrease in NK cell numbers following cytokine-deprivation was more pronounced in NKG2A^+^KIR^+^CD57^-^and.NKG2A^+^KIR^-^CD57^-^ than in more differentiated NKG2A^-^KIR^+^CD57^-/+^ NK cells (Figure 2D).

In summary, these data reveal a subset-dependent susceptibility to IL-15 withdrawal linked to their relative proliferative capacity.

### IL-15 withdrawal results in increased induction of apoptosis

We examined if the observed loss in NK cell numbers was due to an increased rate of apoptosis. To this end, we treated NK cells for 6 days with high-dose IL-15 and analyzed the activation of caspase-3 during the subsequent additional 48 h incubation period in the presence and absence of IL-15. Caspase-3 induction was evident after 48 h post cytokine withdrawal (Figure 3A), which is why we chose to perform all further analyses at this time point. Whereas withdrawal of IL-15 from NK cells pretreated with 10 ng/ml of IL-15 resulted in a significant increase in caspase-3 activation after 48 h, no withdrawal-induced caspase-3 activation was observed in NK cells pretreated with 1 ng/ml of IL-15 (Figure 3B). Stratification of caspase-3 activation by distinguishing between slowly (generation 0-1) and rapidly (generation 2+) cycling NK cells (Figure 3C) revealed a more pronounced caspase-3 activation in rapidly cycling NK cells (Figure 3D).

**Figure 3.**
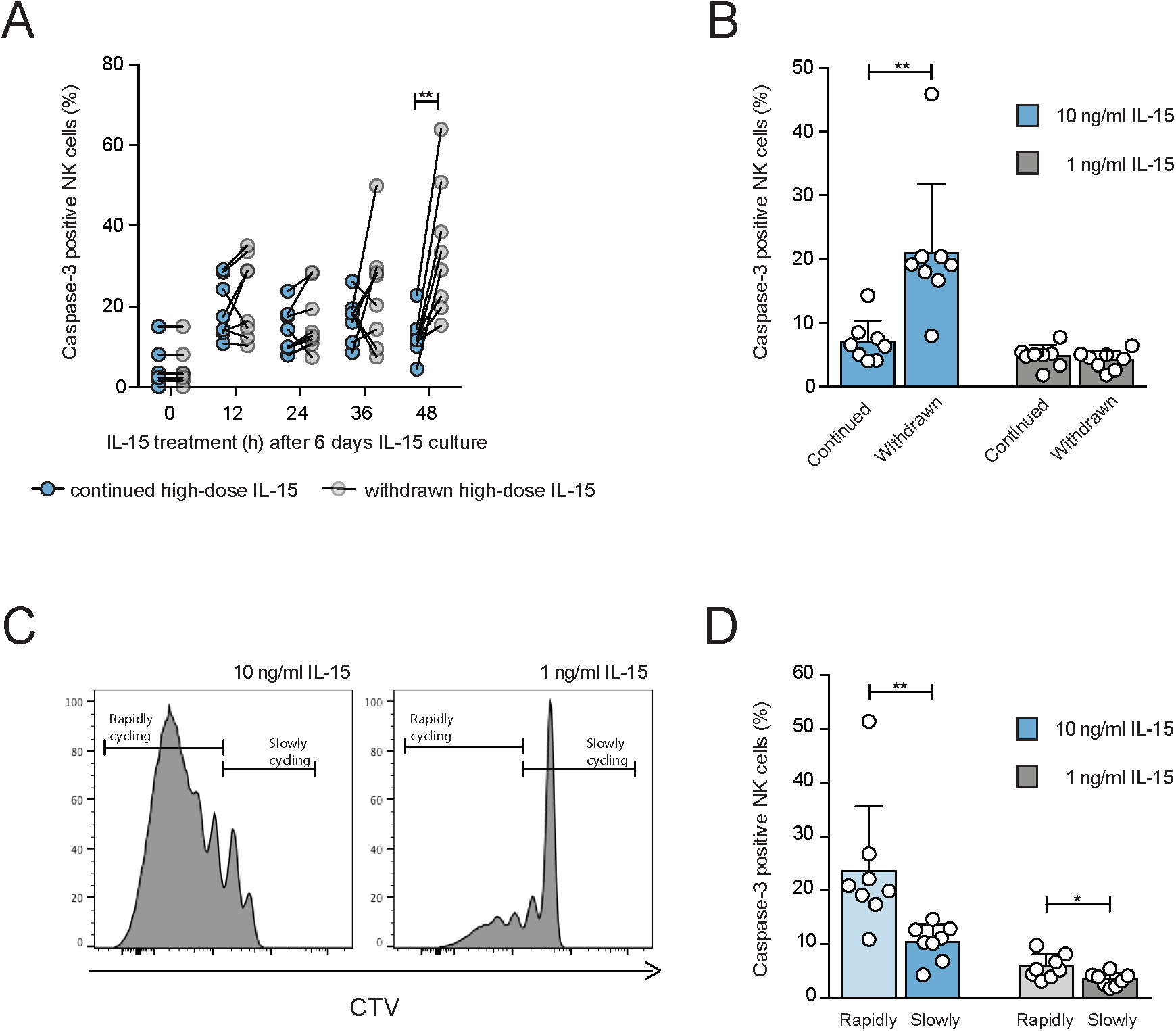
**A)** NK cells were pretreated with high-dose IL-15 for 6 days, harvested and then culture for up to 48 h with or without high-dose IL-15. At the indicated time points NK cells were analyzed for the percentage of active caspase-3 positive cells (n = 8). **B)** NK cells were treated for 6 days with high-(10 ng/ml) or low-dose (1 ng/ml) IL-15, harvested and incubated for additional 48 h with (+) or without (-) the prior IL-15 dose. At the end of the 48 h treatment the percentage of active caspase-3 positive NK cells was measured (n = 8). **C)** Representative histogram of the CTV-dilution in NK cells treated for 6 days with high-(10 ng/ml) or low-dose (1 ng/ml) IL-15. A gate was created around slowly (0-1 cell divisions) and rapidly cycling cells (≥2 cell divisions). **D)** NK cells were treated for 6 days with high-(10) or low-dose (1) IL-15 and incubated for additional 48 h without IL-15. The percentage of active Caspase-3 positive cells was evaluated for rapidly and slowly cycling NK cells (n = 8). Significance were calculated using a Wilcoxon test (p-values: *<0,05; **<0,01).

### Dose-depending up-regulation of BCL-2 family members upon IL-15 treatment

To get an unbiased view on pro-and anti-apoptotic networks following stimulation of NK cells with IL-15 we performed RNA sequencing analysis on NK cells at baseline and after 5 days of IL-15 stimulation (Figure 4A). The most prominent up-regulation was observed for *BIRC5*, a member of the inhibitor of apoptosis (IAP) family. Alongside its anti-apoptotic characteristics, *BIRC5* is mainly involved in the regulation of cell proliferation during chromosomal-microtubule attachment, spindle assembly checkpoint and cytokinesis (29). Among the most significantly up regulated genes observed was BAX, a core member of the intrinsic apoptosis pathway and direct target of BIM (27). However, BCL2, which inhibits the apoptosis pathway at the level of BAX, was also significantly up regulated, potentially balancing the pro-apoptotic axis through BIM/BAX (Figure 4A and Supplementary Table 1).

**Figure 4.**
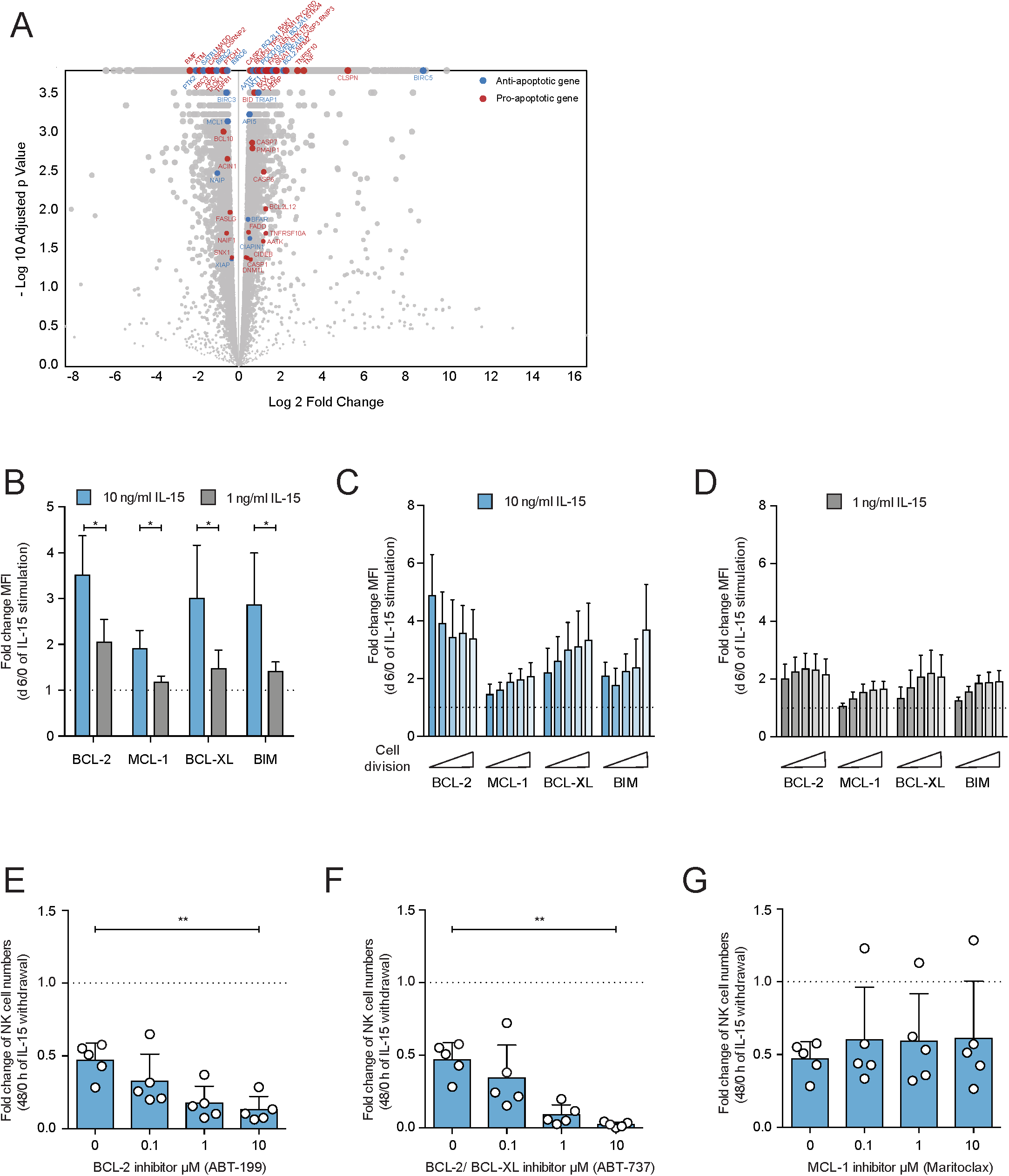
**A)** Global RNA-sequencing analysis comparing baseline to proliferating NK cells after 5 days of IL-15 stimulation. Anti-and pro-apoptotic genes that were significantly differentially expressed (cutoff > 1.5) are highlighted in blue or red, respectively (n=3). **B-D)** NK cells were treated for 6 days with high-(10 ng/ml) or low-dose (1 ng/ml) IL-15 and the fold increase of the expression of different anti-(BCL-2, MCL-1, BCL-XL) and pro-apoptotic (BIM) proteins between day 6 and the beginning of the IL-15 treatment were plotted. The fold increase was calculated for bulk NK cells and for dividing NK cells treated with high-or low-dose IL-15 (n = 7). Significance were calculated using a Wilcoxon test. **E-G)** NK cells were treated for 6 days with high-dose IL-15 and afterwards incubated without IL-15 for additional 48 h in the presence of different concentrations of a BCL-2 (ABT-199; **E**), BCL-2/BCL-XL (ABT-737; **F**) or a MCL-1-inhibitor (Maritoclax; **G**). The fold decrease in NK cell numbers between 48 h after and before IL-15 withdrawal was evaluated (n = 5). Two experiments were excluded from the analysis due to a missing effect of IL-15 withdrawal within the control samples. Significance of the fold decrease in NK cell numbers upon IL-15 withdrawal in the presence of different inhibitors was calculated between the individual inhibitor concentrations and the control using a Friedman test (p-values: * <0,05; ** <0,01).

Next, we analyzed the protein expression of anti-apoptotic BCL-2, MCL-1 and BCL-XL as well as pro-apoptotic BIM in primary human NK cells after 6 days of IL-15 treatment. Upon IL-15 treatment NK cells up-regulated all three anti-apoptotic proteins, but also BIM (Figure 4B). All four BCL-2 family members were significantly more up regulated in cells treated with high-dose IL-15 compared to low-dose IL-15. Interestingly, whereas MCL-1, BCL-XL and BIM expression increased with the number of cell divisions, BCL-2 expression was highest in slowly or non-dividing NK cells after incubation with high-dose IL-15 (Figure 4C-D). Next, we incubated high-dose IL-15 pretreated NK cells for 48h without IL-15 in the presence of different concentrations of BCL-2 (ABT-199) and BCL-2/BCl-XL (ABT-737) inhibitors (Figure 4E-G). Whereas BCL-2 inhibition led to a negative influence on NK cell survival during IL-15 withdrawal, MCL-1 inhibition with Maritocax had no effect, even at high concentrations. Notably, as a control, Maritoclax decreased MCL-1 levels and survival in K562 cells (Supplementary Figure 2A-B), but had only modest effects on MCL-1 levels in NK cells (Supplementary Figure 2C). These data indicate a crucial role for BCL-2 in protecting NK cells during IL-15 withdrawal and suggest that this protection is more fragile in cells that have undergone multiple rounds of cell division.

### Altered BCL-2/BIM ratio following IL-15 withdrawal

Based on the observation that IL-15 induced a profound and dose-dependent up-regulation of both pro-and anti-apoptotic proteins, we addressed whether the susceptibility to IL-15 withdrawal in proliferating NK cells was linked to a selective decrease in anti-apoptotic proteins. We found that the expression of all three anti-apoptotic proteins decreased following 48 h of culture in the absence of IL-15 (Figure 5A). The effect was more pronounced in NK cells pretreated with high-dose IL-15. Intriguingly, the expression of pro-apoptotic BIM also decreased 48h after IL-15 withdrawal, although not as prominent as BCL-2, leading to altered BCL-2/BIM ratios (Figure 5B). The balance between anti-and pro-apoptotic proteins is critical for the induction of apoptosis (30). We found that the BCL-2/BIM ratio was significantly lower in rapidly dividing NK cells upon IL-15 stimulation and decreased further upon IL-15 withdrawal, dropping to levels below those in resting NK cells (Figure 5C). Thus, the altered balance between BCL-2 and BIM in highly proliferating NK cells may contribute to the NK cell death observed after cytokine withdrawal.

**Figure 5.**
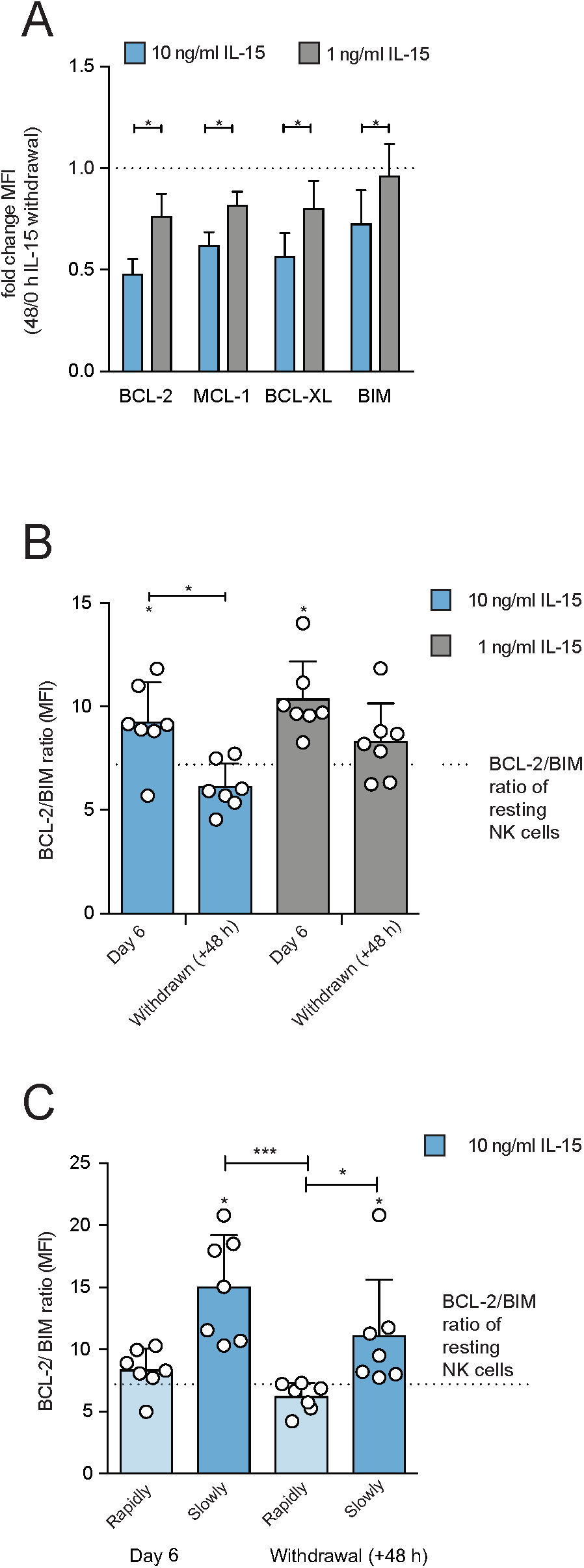
**A)** NK cells were treated for 6 days with high-(10 ng/ml) or low-dose (1 ng/ml) IL-15 and then incubated for additional 48 h without IL-15. The fold decrease of the expression of different anti-(BCL-2, MCL-1, BCL-XL) and pro-apoptotic (BIM) proteins after 48 h compared to the beginning of IL-15 withdrawal was plotted for bulk NK cells (n = 7). Significance was calculated using a Wilcoxon test. **B-C)** The BCL-2/BIM ratio was calculated for freshly isolated NK cells (dotted line) and for NK cells after 6 days of high-(10 ng/ml) or low-dose (1 ng/ml) IL-15 treatment or after 48 h of IL-15 withdrawal. Results were plotted for bulk (**B**) and for rapidly vs. slowly cycling (**C**) NK cells (n = 7). The increase or decrease of the BCL-2/BIM ratio compared to resting NK cells (**B**-**C**), was calculated using a Wilcoxon signed rank test (stars alone), while differences within the ratio between differently treated NK cells was calculated using a Friedman test (stars on line).

### Treatment with high-dose IL-15 induces the expression of the BIM S splice variant

Although the ratio BCL-2/BIM ratio in proliferating NK cells decreased to levels below baseline, the effect of IL-15 withdrawal was rather subtle (Figure 5C). Therefore, we next investigated the potential role of different BIM splice variants in the enhanced susceptibility of highly activated NK cells to apoptosis. There are at least 18 known BIM splice variants with various potential to induce apoptosis, of which BIM extra long (BIM EL), BIM long (BIM L) and BIM S are the major ones (31). In particular, BIM S has been shown to be more potent in inducing apoptosis (32). Treatment with IL-15 led to an increase in all three BIM splice variants over 6 days, with BIM S demonstrating the strongest up-regulation, in particular in NK cells exposed to higher concentrations of IL-15 (Figure 6A-B). To further evaluate the influence of proliferation, we monitored the three splice variants in FACS sorted proliferating and non-proliferating NK cells after 4 days of IL-15 stimulation using the Peggy Sue instrument for protein analysis of small sample volumes (33). We found that proliferating NK cells had a higher expression of the BIM S variant than non-proliferating NK cells (Figure 6C). In contrast, no clear differences were observed for the other two BIM variants when stratifying for the degree of proliferation. Corroborating the flow cytometry data in Figure 4C, we found that BCL-2 expression was lower in proliferating than in non-proliferating NK cells when treated with high-dose IL-15 (Figure 6C). We then followed the expression of the splice variants during IL-15 withdrawal and found that BIM S levels remained high for 24 h before dropping to marginal levels (Figure 6D and Supplementary Figure 3A). This was paralleled with the amount of BCL-2 roughly halving by 24 h, mirroring the flow cytometry data in Figure 5A. Together, these data suggest that the unique sensitivity of proliferating NK cells to apoptosis is linked to a selective accumulation of the toxic BIM S splice variant together with diminishing levels of BCL-2.

**Figure 6.**
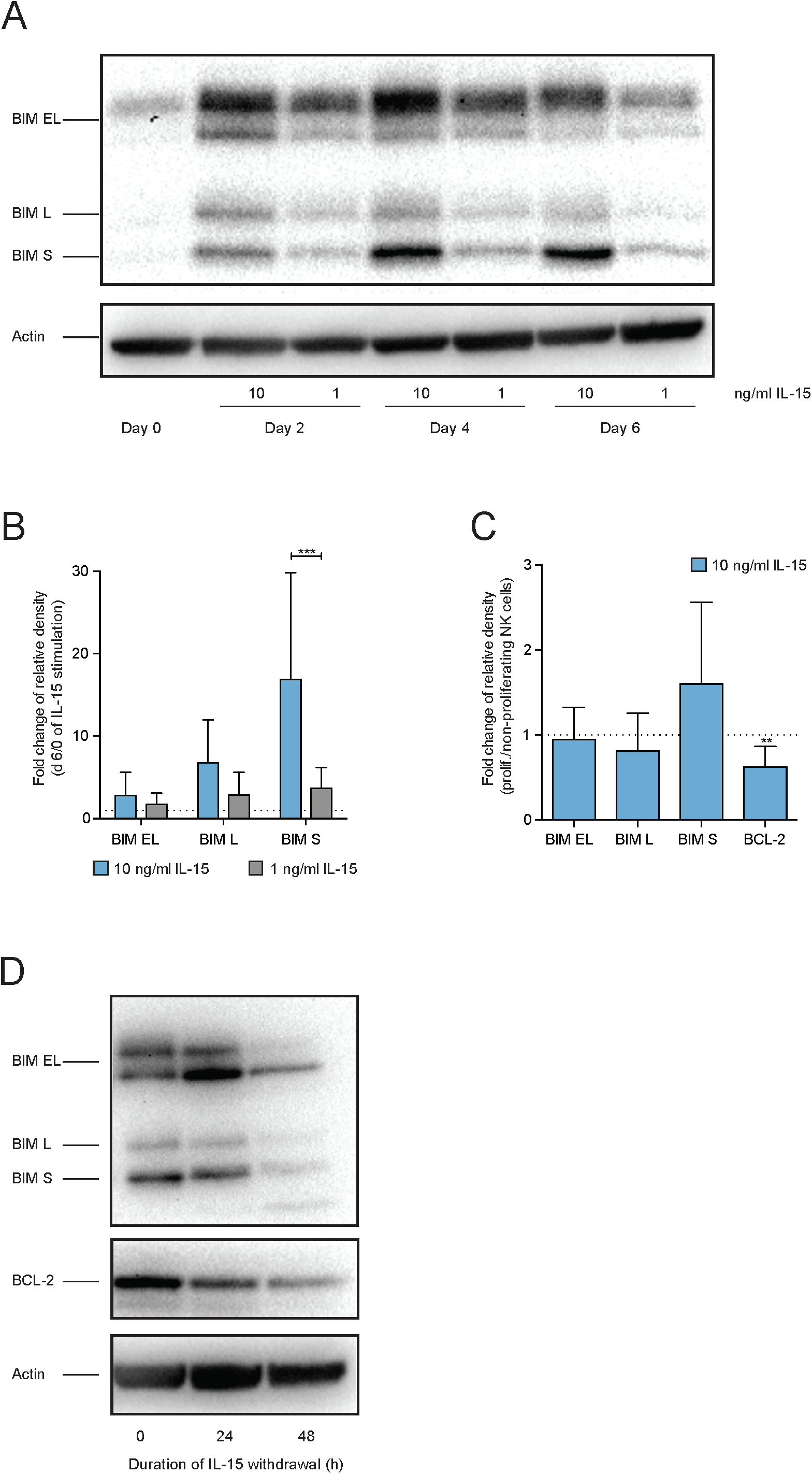
**A)** A representative western blot analysis for the expression of BIM splice variants in freshly isolated NK cells (day 0) and NK cell treated for 2, 4 or 6 days with high-(10) or low-dose (1) IL-15. Actin was used as a loading control. **B)** NK cells were treated with high-(10 ng/ml) or low-dose (1 ng/ml) IL-15 for 6 days and the fold increase of the expression of different BIM splice variants after 6 days of IL-15 treatment compared to resting NK cells was evaluated (n = 6). **C)** NK cells were treated with high-dose IL-15 for 4 days, FACS-sorted based on their CFSE dilution into proliferating (at least one prior cell division) or non-proliferating (no cell division) NK cells and then analyzed at a Peggy Sue^™^ machine. The fold change between the expressions of BIM splice variants or BCL-2 in non-proliferating vs. proliferating NK cells was calculated (n = 10). **D)** A representative western blot analysis for the expression of BIM splice variants and BCL-2 after 0, 24 and 48 h of IL-15 withdrawal in NK cell pretreated with high-dose (10) IL-15 for 6 days. Actin was used as a loading control. Significance between NK cells treated with high-(10 ng/ml) or low-dose (1 ng/ml) IL-15 for 6 days in the expression of the different BIM splice variants was calculated using a Wilcoxon test (**B**). Significance between non-and proliferating NK cells (**C**) for their expression of BIM splice variants and BCL-2 was calculated using Wilcoxon signed rank test (p-values: ** <0,01; ***<0,001).

## Discussion

Understanding how cytokine-priming influences NK cell homeostasis *in vivo* and *in vitro* is essential for the development of NK-cell based immunotherapies. The transfer of highly activated *in vitro* expanded NK cells into a lymphopenic, pre-conditioned host represents a critical phase due to the sudden change in cytokine concentration. The degree to which the transfer itself leads to loss of function and cell death of donor-derived NK cells is likely to depend on many factors. This includes the length and extent of prior activation, the combination of cytokines used, and the application/implementation of supportive systemic cytokine treatment regimes in the patient. In this study we established a robust platform to study the fate of discrete NK cell subsets following abrupt withdrawal of IL-15, mimicking a scenario where IL-15 activated and expanded NK cells are transferred into a patient.

During the course of viral infections, murine NK cells undergo expansion followed by a contraction phase due to reduction in cytokine levels when the infection resolves. Members of the BCL-2 family play a crucial role in regulating the fate of immune cells during this process (34). We observed a dose-dependent and subset-specific up-regulation of pro-and anti-apoptotic BCL-2 family members during IL-15 stimulation followed by a down-regulation upon IL-15 withdrawal. These results corroborate previous reports in human (35) and murine NK cells (24) demonstrating that IL-15 stimulation up-regulates anti-apoptotic proteins such as BCL-2 and MCL-1 in NK cells. In addition, we show that IL-15 withdrawal results in a decrease of BCL-2 and MCL-1 in primary human NK cells, which has so far only been described in murine NK cells (24). In human NK cells, up-regulation of BCL-2 during a short stimulation with IL-15 was dependent on STAT5, but not mTOR, and was linked to improved survival upon cytokine withdrawal *in vitro* and *in vivo* (20). In murine T cells it has been reported that long-term surviving effector CD8^+^ T cells had higher anti-apoptotic Bcl-2 protein levels compared to their short-lived counterparts, along with higher levels of pro-apoptotic Bim (36). The increased Bcl-2 protein levels enabled murine effector CD8^+^ T cells to tolerate the higher Bim levels. *In vivo* administration of IL-7 or IL-15 into C57BL/6 mice infected with LCMV increased Bim protein levels within effector CD8^+^ T cells, while inhibition of Bcl-2 resulted in decreased Bim levels. These results suggested that Bcl-2 and Bim expression is coordinated in murine T cells and that the Bcl-2 protein levels determinate the amount of Bim protein levels a cell is able to tolerate. *Ex vivo* stimulation of murine NK cells with IL-15 was found to down-regulate the expression of Bim via transcriptional repression and increased proteasomal degradation (24). Upon IL-15 withdrawal, Bim expression increased, leading to cell death in murine NK cells. Furthermore, several groups demonstrated that changes in the Bcl-2/Bim ratio can render murine T and NK cells sensitive to cell death (24, 36, 37). Specifically, studies in conditional knock-out mice suggest that BCL-2 is a non-redundant survival protein for murine NK cells at rest, whereas MCL-1 is the dominant survival protein during proliferation (38). The importance of the BCL-2/BIM dynamics is supported by our findings in primary human NK cells, since we observed a proliferation-dependent loss of BCL-2 during IL-15 withdrawal, resulting in low BCL-2/BIM ratios and increased NK cell death. While increased levels of MCL-1 and increased BCL-2/BIM ratios appear to protect from apoptosis during ongoing proliferation, such highly “addicted” NK cells show a drop in BCL-2/BIM ratios below the level of resting NK cells after IL-15 withdrawal. Surprisingly, pharmacological inhibition of MCL-1 with Maritoclax showed minimal effect on survival in our *in vitro* model whereas BCL-2 inhibition led to a near complete cell death during IL-15 withdrawal.

An additional factor contributing to the susceptibility of rapidly proliferating NK cells to IL-15 withdrawal was their differential expression of BIM splice isoforms during IL-15 stimulation. The BIM protein is known to have at least 18 different splice variants with different abilities to induce apoptosis (31). The activity of the two variants BIM EL and L to induce apoptosis is regulated via phosphorylation at several phosphorylation sites (39). In addition, both isoforms contain the dynein light (L) chain-binding domain (DBD), enabling them to be sequestered by dynein on microtubules (40). In response to apoptotic stimuli they are released from the microtubules and become activated. In contrast, the BIM S splice variant is neither controlled by posttranscriptional phosphorylation nor by binding to the microtubules. Furthermore, it is able to directly bind BAX making it the most potent apoptosis inducing BIM isoform (31). Here, we observed that IL-15 stimulation in general increased the expression of all three major BIM variants, whereas high-dose IL-15 preferentially up-regulated the highly cytotoxic BIM S variant, particularly in cells that had undergone extensive cell division. One explanation for this phenomenon could be an increased production of reactive oxygen species (ROS) in proliferating NK cells. It is known that upon murine NK cell expansion during MCMV infection, proliferating NK cells accumulate increased levels of reactive oxidative species (ROS) due to the accumulation of depolarized mitochondria. Removal of damaged mitochondria/ROS via mitophagy was crucial for the survival of NK cells in this model (41). ROS can influence the expression levels of BCL-2 and BIM. In T cells, it has been shown that *in vivo* activation with staphylococcal enterotoxin B (SEB) leads to decrease Bcl-2 levels in stimulated T cells and subsequently to cell death. Bcl-2 levels could be restored by blocking ROS production using the antioxidant Mn(III)tetrakis(5,10,15,20-benzoic acid)porphyrin (MnTBAP) (30). In addition, it is known that increased ROS levels are able to stimulate BIM expression via the ROS-JNK-BIM pathway (42).

A recent study described the impact of distinct dose scheduling of high-dose IL-15 stimulation on NK cells. Continuous treatment with high-dose IL-15 for 9 days was more potent at inducing robust NK cell proliferation than IL-15 treatment for 9 days with a three-day lack of IL-15 (day 4-6). However, intermitting IL-15 treatment resulted in improved survival and function of NK cells compared to continuous IL-15 treatment, which was linked to the induction of a cell cycle arrest genes and reduced mitochondrial respiration due to mTOR signaling (43). Interestingly, cell cycle arrest genes, including GADD45 and P53 are known to positively regulate the expression of BIM, whereas P53 has a negative impact on BCL-2 expression levels (44, 45). In human NK cells, mTOR signaling is tightly associated with proliferation (Pfefferle et al., unpublished observation). Chronic AKT signaling, which is a downstream target of mTOR signaling, leads to down-regulation of BCL-2 in T cells (46), which aligns with the observation of a gradual decline in BCL-2 when cells undergo cell division, as noted in the present study. On the contrary, mTOR signaling suppresses the expression of BIM, since inhibition of mTOR signaling up-regulates BIM expression in various cell types (47, 48). This differs from the observation in primary NK cells as we noted a robust increase in BIM levels and in particular of the BIM S splice variant in proliferating NK cells. Therefore, potential differences in mTOR signaling cannot fully explain the increased susceptibility of high-dose IL-15 treated NK cells towards IL-15 withdrawal.

In summary, our results reveal a subset and proliferation-dependent alteration in BCL-2/BIM ratios with increased levels of the toxic BIM S splice variant. Together these changes sensitize IL-15 addicted NK cells to cytokine withdrawal leading to apoptosis. These insights may pave the way for gene-editing approaches aiming to restore BCL-2/BIM ratios or interfering with BIM S or its downstream targets prior to adoptive transfer to prolong the *in vivo* persistence of expanded NK cell products. Alternatively, it may be possible to rescue the addicted phenotype by titrating the level of IL-15 before *in vivo* infusion of the final cell therapy product.

## Acknowledgments

We would like to thank Drs Johannes Landskron and June H. Myklebust for their kind assistance in establishing the fluorescent bar coding technology and Dr Jai Rautala for critical reading of the manuscript.

